# Transcriptional regulation by Tbx2 paralogs creates diversity within photoreceptor subtypes

**DOI:** 10.64898/2026.07.20.739565

**Authors:** Carinna M. Householder, Autumn S. Lee, Sofia Apgar, Annabella K. Rinaldi, Juan Angueyra

**Affiliations:** Department of Biological Sciences, University of Maryland, 4094 Campus Dr, College Park, MD, 20742, USA; Brain and Behavior Institute, University of Maryland, 4094 Campus Dr, College Park, MD, 20742, USA

**Keywords:** Vision, retina development, transcription factor, subfunctionalization, photoreceptors, cell subtypes, cell fate

## Abstract

Photoreceptors are retinal neurons whose subtype-specific opsin expression and mosaic organization enable parallel encoding of spectral and spatial visual information. Yet, the transcriptional mechanisms establishing and maintaining subtype identity remain poorly understood. We investigated the role of the T-box transcription factors *tbx2a* and *tbx2b* in zebrafish photoreceptor development and found evidence for paralog subfunctionalization across photoreceptor lineages. Tbx2b is required for UV-cone generation and prevents fate switching toward rods, while Tbx2a is constrained to UV-cone generation without involving rods. These factors also maintain cone identity by repressing inappropriate M-cone fate. Despite major changes in photoreceptor composition, *tbx2* mutants retain luminance- and motion-driven vision, whereas prey capture is abolished only in *tbx2b* mutants and not *tbx2a* mutants. Our findings identify the *tbx2* paralogs as key regulators of cone-subtype generation and maintenance and reveal how developmental programs diversify photoreceptor fate to appropriately support visually-guided behaviors important for survival.

## INTRODUCTION

As organs develop, progenitors generate specialized cell types that fulfill specific functions, by progressively changing gene expression. Transcription factors directly orchestrate these transitions by binding to regions of regulatory DNA to activate or repress target genes, establishing the transcriptional differences that eventually define cellular identity.^1,2^ Because many developmental decisions involve closely related alternative fates, these regulatory programs must be implemented with high precision and robustness to ensure both correct fate generation and long-term maintenance of subtype identity.^3^ Understanding how transcription factors achieve this balance remains a central challenge in developmental biology. Photoreceptors, the cells that transform light information into electro-chemical signals and relay them into retinal circuits, can be divided into distinct subtypes, which differ in morphology, gene expression, density and wiring.^4,5^ Zebrafish possess five photoreceptor subtypes, rods and four cones each spectrally sensitive to long (L), medium (M), short (S), and ultraviolet (UV) wavelengths of light. All of the photoreceptor subtypes are generated from regulatory programs that are conserved across vertebrates.^6^ Functionally and molecularly, zebrafish L cones (expressing *opn1lw* paralogs) are evolutionarily related to human L and M cones (expressing *OPN1LW* and *OPN1MW*), and zebrafish UV cones (expressing *opn1sw1*) to the human S cones (expressing *OPN1SW1*).^7–9^ Zebrafish S (expressing *opn1sw2*) and M cones (expressing *opn1mw* paralogs) are absent in humans, as these subtypes were lost in mammals.^10^ The differences in spectral sensitivity of photoreceptor subtypes are mediated by the expression of specific opsins, the receptors that initiate phototransduction.^11^ Although photoreceptor identity is closely linked to opsin expression, their subtype-specific differences extend beyond that to transcription factors and their subsequent regulatory networks during development.^10,12,13^ In addition, photoreceptor subtypes in zebrafish are arranged in a highly stereotyped mosaic that efficiently samples visual information. Establishing such ordered architecture requires producing the correct number of each subtype and arranging them in a defined pattern, which also is under transcriptional regulation.^14,15^ Perturbations in photoreceptor composition compromise visual function and cause disease phenotypes. For example, in humans, mutations in the transcription factor NR2E3 cause the “*enhanced S-cone syndrome*”: by disrupting the transcriptional programs used to generate rods, rod progenitors are rerouted to become S cones, leading to severe visual defects that include night blindness.^16^ This study aims to understand how transcriptional programs are initiated and maintained to generate the five photoreceptor identities of vertebrates.

During development, photoreceptors are derived from a pool of proliferative retinal progenitors that undergo progressive fate restriction and differentiation. Commitment towards photoreceptor identity requires activating genes for photoreceptors while repressing genes associated with the other cellular identities of the retina (*e.g.*, bipolar or horizontal cells).^17–20^ Although we know a great deal about the transcription factors that govern rod development,^21,22^ the transcriptional logic that specifies and stabilizes cone subtype identity is less well resolved, and members of the T-box family have recently emerged as candidates for cone-fate regulation.^12,23^ In zebrafish, loss of *tbx2b* eliminates UV cones and is accompanied by a near-matched increase in rods, consistent with a fate switch in photoreceptor precursors.^24,25^ However, how T-box factors control cone subtype identity more broadly—and whether they act primarily in generation versus maintenance of subtype-restricted gene expression programs—remains unclear. Although the role of *tbx2b* in generating UV cones is well described, its paralog *tbx2a*, is poorly characterized, despite subtype-enriched expression patterns in which *tbx2b* is expressed by UV and S cones, while *tbx2a* is expressed by UV and L cones.^12^ Here, we define how *tbx2b* and *tbx2a* have become subfunctionalized to control cone-subtype generation and maintenance, revealing distinct paralog functions that safeguard UV-cone identity and prevent inappropriate activation of M cone gene-expression programs in S and L cones. Moreover, we use behavioral assays to discern the impact of such changes in photoreceptor subtypes for vision.

## RESULTS

Photoreceptor subtypes arise from shared progenitors and remain closely related at the molecular level, raising a key developmental question: what transcriptional mechanisms separate subtype-specific fates during development and are they involved in preserving and maintaining these fates?^26^ We use zebrafish to investigate how the duplicated T-box transcription factors, Tbx2b and Tbx2a, have subfunctionalized to initiate and maintain regulatory networks important for cone and rod lineages. The results are organized to connect mechanism to function. First, we establish that the Tbx2 paralogs are not equivalent for the generation of UV cones and rods: *tbx2b* mutants cannot form UV cones and instead form rods, whereas *tbx2a* mutants have significant loss of UV cones but rods are unaffected. Second, we find that both Tbx2b and Tbx2a have shared roles to maintain two of the other cone-subtype identities: *tbx2b* mutants ectopically express M opsin in S cones and *tbx2a* mutants ectopically express M opsin in L cones. Furthermore, we link how these subtype-specific defects impact vision in *tbx2* mutants. We find that luminance and motion processing are preserved despite the multiple alterations to photoreceptor subtypes. In addition, while complete loss of UV cones in *tbx2b* mutants abolishes prey capture, remaining UV cones in *tbx2a* mutants are functional and able to support hunting.

### Tbx2a and Tbx2b are essential but not equivalent for UV-cone generation

Previous studies have characterized mutations in the coding or regulatory regions of *tbx2b* that produce the *fby (from beyond)* and *lor (lots of rods)* phenotypes respectively, where UV cones are not generated during development and rod densities increase, suggesting a switch in photoreceptor fate.^24,25^ To characterize the role of Tbx2b and Tbx2a in the generation of UV cones and rods, we used CRISPR-based mutagenesis to create germline mutants for both *tbx2* genes (*tbx2b*^mcp1^ and *tbx2a*^mcp2^ mutants), targeting the third exon that encodes the highly-conserved amino-acid residues that bind directly to DNA and are critical for the gene-regulatory functions of these transcription factors (Figure 1A-1F).^27^ We quantified UV-cone and rod densities and homotypic nearest-neighbor distances (NND) in the central retina of 5 days post fertilization (dpf) larvae using transgenic reporter lines for UV opsin (*opn1sw1:nfsB-mCherry*) and rhodopsin (*xOPS*:GFP) (see STAR Methods). In wild-type larvae, UV cones are abundant and evenly spaced, while rods are sparse at these young ages (Figure 1G). We find that, in agreement with previous studies, *tbx2b* mutants have virtually no UV cones and a marked increase in rod densities and decreased NND between rods (Figure 1H; 1J-L); we named this phenotype *nuri (no uv rods increase*) which phenocopies the previously described *fby* phenotype.^24,25^ The absence of UV cones disrupts the organization of the cone mosaic, evidenced by misalignment of the non-UV cone subtypes in the larval-lattice (Figure 1J-1K).^28^ In contrast, *tbx2a* mutants have a marked yet incomplete loss of UV cones (∼36% mean reduction in UV cones; Figure 1I) and a corresponding increase in NND between UV cones but without any changes in rod densities or their spacing (Figure 1J-L); we named this phenotype *nalou* (*not a lot of UV).* Even with the presence of this small fraction of UV cones, the cone mosaic remains altered and the distribution of UV cones is discontinuous (Figure 1I). To confirm that this *nalou* phenotype is not a result of residual protein function, we performed a genetic complementation test. We induced an independent mutation in the first exon of *tbx2a* (*tbx2a*^mcp3^ mutants) and generated crosses with *tbx2a*^mcp2^ mutants (Figure S1A-D). We could not identify differences in UV cone or rod densities between *tbx2a*^mcp3^ mutants, *tbx2a*^mcp2^ mutants, or *tbx2a^mcp2^/tbx2a^mcp3^*double heterozygous mutants (Figure S1E-L). Our genetic complementation results suggest that all mutations we generated in *tbx2a* cause a complete loss of function and cause the *nalou* phenotype.

**Figure 1:**
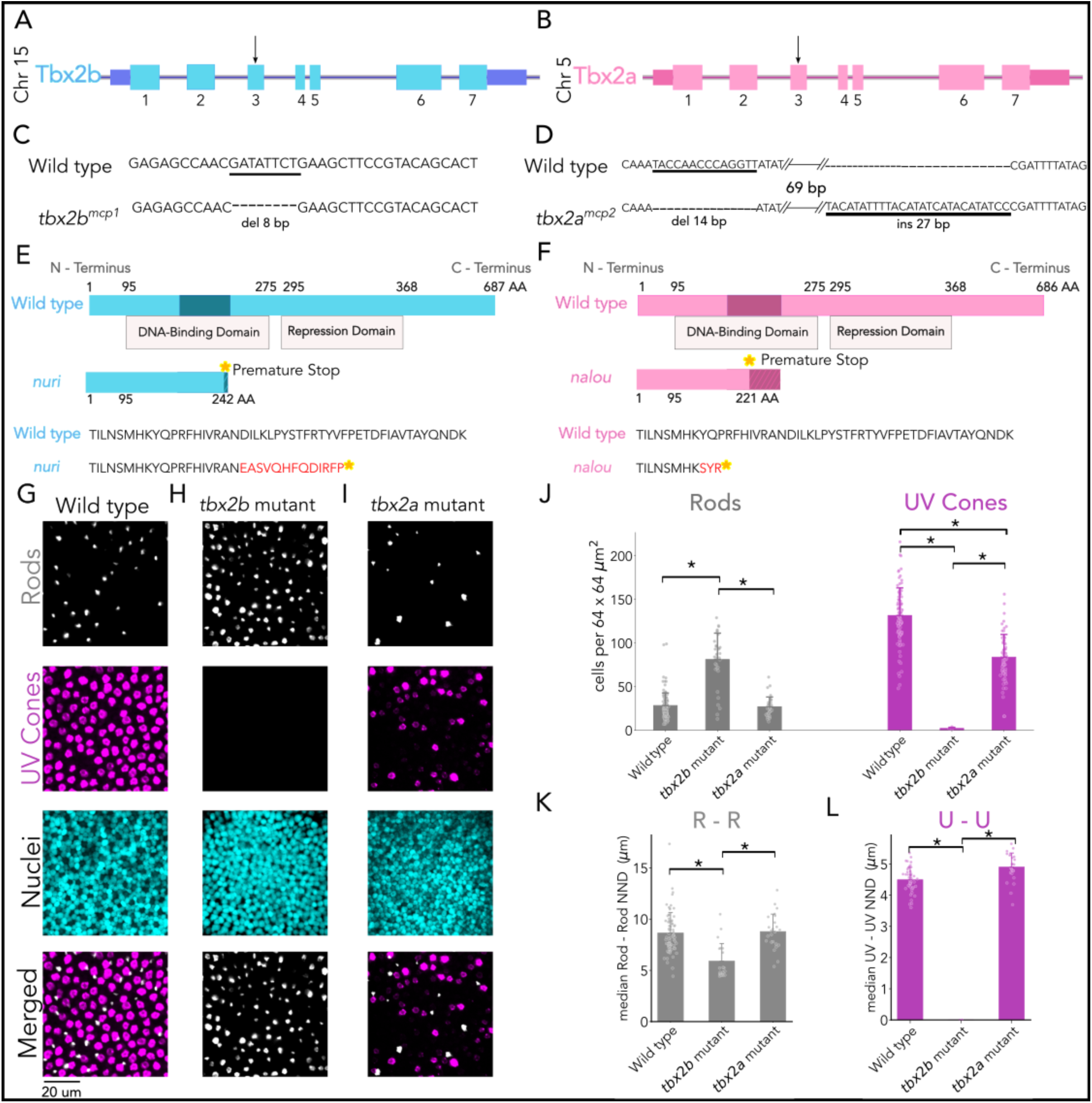
*tbx2b* and *tbx2a* are essential for UV cone generation but not equivalent. **(A-B)** Map of *danio rerio tbx2b* (blue) and *tbx2a* (pink) genes on chromosomes 15 and 5. Both coding regions span 5.7 kB consisting of seven coding exons (thick boxes), six introns (solid lines), and 1.4 kB of 5’ and 3’ UTRs (thin boxes). (**C-D**) Isolated mutations for wild type (top) and mutant (bottom) alleles. In *tbx2b* mutants, a 8 bp deletion (del) disrupts exon 3. In *tbx2a* mutants, a 14 bp deletion (del) and 27 bp insertion (ins) disrupts exon 3, encoding the DNA-binding domain. (**E-F**) Protein diagram of Tbx2b and Tbx2a mutations. Predicted primary sequence of proteins for wild type (top) and mutants (bottom). The amino-acid positions that result from exon 3 are shaded in darker shades for each protein. The highly conserved DNA-binding domain and the T-box repression domain are denoted by the shaded boxes underneath. Both mutations result in frameshifts that introduce a premature stop codon (indicated by yellow star). (**G-I**) Representative confocal images of the central retina of wild type (left), *tbx2b* mutant (center), and *tbx2a* mutant (right) at 5 dpf, in double transgenic larvae with labeled rods (grey) and UV cones (magenta). (**J**) Quantification of rod and UV-cone densities in wild-type, *tbx2b* and *tbx2a* mutant larvae. Compared to controls, *tbx2b* mutants have a significant increase in rods while *tbx2a* mutants do not; both *tbx2* mutants have a marked decrease in UV cones (Kruskal-Wallis H_Rods_=50.859, p_Rods_=9.03 × 10^−13^, n*_wild type_* = 120, n*_tbx2b_*=34, n*_tbx2a_*=34; Conover-Iman posthoc corrected p-values: control vs. *tbx2b* p=2.51 × 10^−13^, *tbx2b* vs. *tbx2a* p=1.85, × 10^−10^; Kruskal-Wallis H_UV_=66.402, p_UV_=3.81 × 10^−14^, n*_wild type_* = 87, n*_tbx2b_*=3, n*_tbx2a_*=61; Conover-Iman posthoc corrected p-values: control vs. *tbx2b* p=3.12 × 10^−6^, control vs. *tbx2a* p=4.27 × 10^−18^). (**K-L**) Quantification of rod and UV cone spacing using median nearest-neighbor distances between homotypic photoreceptor subtypes. *tbx2b* mutants have a significant decrease in the spacing of rods compared to wild type, (Kruskal-Wallis H=26.248, p=1.024 × 10^−6^, n*_wild type_* = 74, n*_tbx2b_*=424, n*_tbx2a_*=20; Conover-Iman posthoc corrected p-values: control vs. *tbx2b* p=1.02 × 10^−6^, *tbx2a vs. tbx2b* p=5.05 × 10^−6^), while *tbx2a* mutant UV cone and rod spacing is not significantly altered (wild type vs. *tbx2a* mutant, p=1.0). Bars represent averages, error bars correspond to standard deviations, and markers correspond to individual retinas.

In summary, our results demonstrate that *tbx2* paralogs in zebrafish have subfunctionalized, in spite of their co-expression in UV cones: *tbx2b* is necessary to generate UV cones by inhibiting rod fate while *tbx2a* is partially dispensable for UV-cone generation and does not influence rod development. Since *tbx2b* is also expressed in S cones and *tbx2a* in L cones,^12^ we decided to explore the development of the other cone subtypes in our *tbx2* mutants.

### Tbx2b regulates M-opsin expression in S cones

To characterize the role of Tbx2b in the development of S cones, we used *tbx2b*^mcp1^ and *tbx2a*^mcp2^ mutants in combination with reporter lines for S opsin (*opn1sw2*:nfsb-mCherry) and M opsin (*opn1mw2*:GFP) and quantified photoreceptor densities and homotypic NND (see STAR methods Key Resources Table). In wild-type larvae, M- and S-opsin expressing cells are regularly distributed with no overlap in expression between reporters—except a small fraction of S cones that express GFP, as described in the original publication of the M-opsin reporter line (Figure 2A).^29^ In both *tbx2b* and *tbx2a* mutants, we find that S cones (mCherry positive) are present in normal densities and their spacing is indistinguishable from wild type, while cells that express M-opsin (GFP positive) have increased density and appear to be spaced irregularly (Figure 2B-E). In *tbx2b* mutants, these supernumerary GFP-positive cells correspond to S cones (mCherry and GFP positive), a sign of ectopic expression of M opsin in S cones. In contrast, S cones in *tbx2a* mutants are not GFP positive and are likely normal, in agreement with the lack of expression of *tbx2a* in S cones (Figure 2D-F). By excluding the mCherry and GFP positive S cones in *tbx2b* mutants, we quantified the density and spacing of GFP-only cells, which we presumed correspond to M cones, and found that their density is not significantly different from wild type animals. Yet, we find a decrease in their nearest-neighbor distance, suggesting that M-cone arrangement but not number is impacted by the absence of UV cones (Figure 2G-H). These results suggest that, in *tbx2b* mutants, M cones are likely normal, but S cones lose the ability to repress M-cone fate, which erodes their subtype-specific identity.

**Figure 2:**
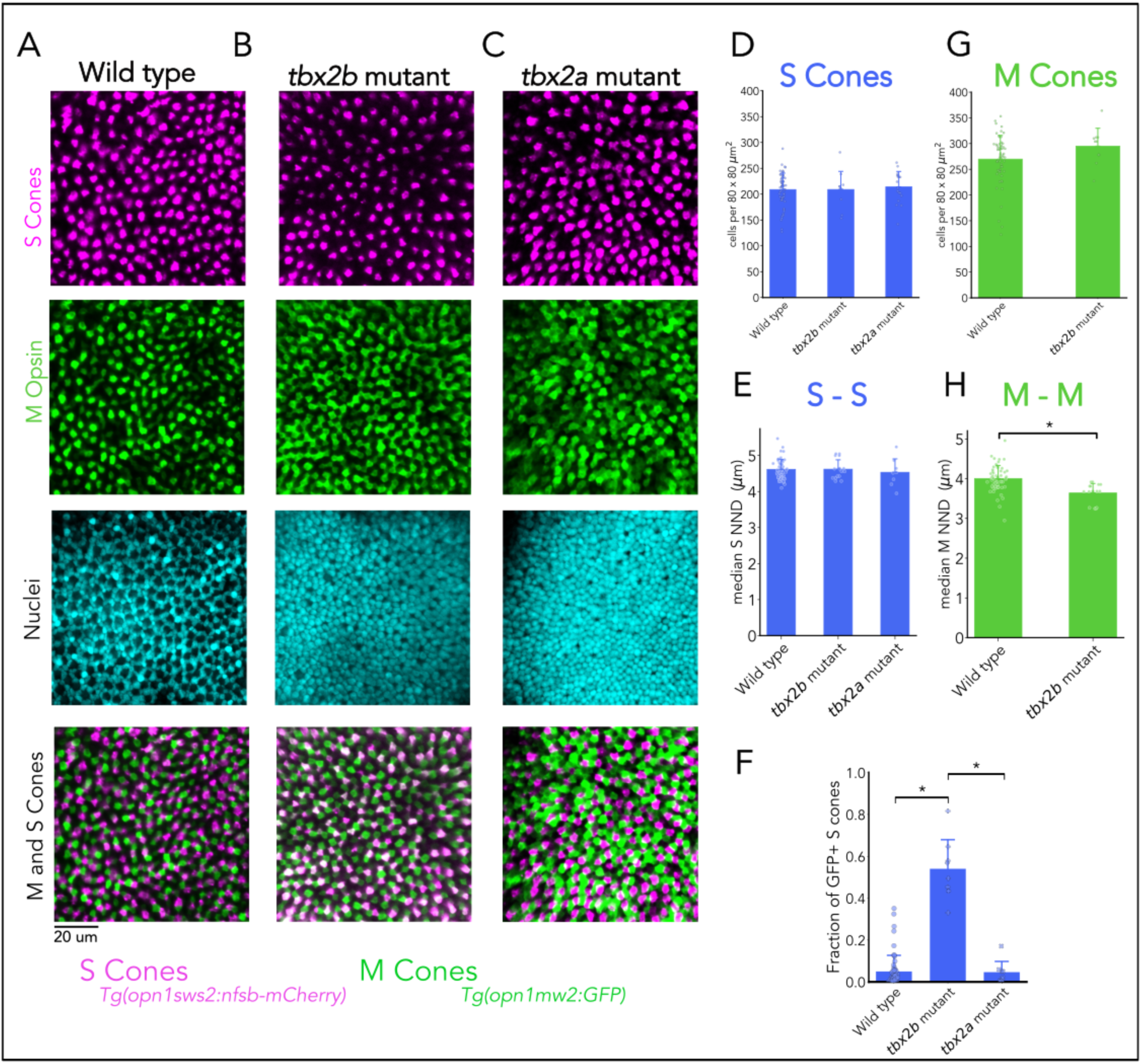
*tbx2b* represses M opsin in S cones. (**A-C**) Representative confocal images of the central retina of wild type (left), *tbx2b* mutants (center), and *tbx2a* mutants (right) at 5 dpf, in double transgenic larvae with labeled M cones (green), S cones (magenta), and nuclei (cyan). Both *tbx2b* and *tbx2a* mutants display an increase in GFP-positive cells. In *tbx2b* mutants the increase in GFP signal is restricted to mCherry-positive cells, whereas *tbx2a* mutants have an increase in GFP signal that does not overlap with mCherry-positive cells. (**D-E**) Quantification of M- and S-cone numbers in wild-type, *tbx2a* and *tbx2b* mutant larvae. Both S- and M-cone numbers showed no significant changes in either *tbx2b* or *tbx2a* mutants compared to wild type. (Kruskal-Wallis H_S_=0.447, p_S_=0.799, n*_wild type_* = 69, n*_tbx2b_*=10, n*_tbx2a_*=18; Kruskal-Wallis H_M_=1.522, p_M_=0.4671). (**F-G**) Quantification of M- and S-cone spacing, using median nearest-neighbor distances. Neither *tbx2b* or *tbx2a* mutants show significant changes in S-cone spacing compared to wild type, while both *tbx2b* mutants show a significant decrease in space between M cones compared to wild type (Kruskal-Wallis H_S_=0.547, p=1.0, n*_wild type_* = 58, n*_tbx2b_*=8, n*_tbx2a_*=9; Kruskal-Wallis H_M_ =16.652, p=2.4 × 10^−4;^ ; n*_wild type_* = 56, n*_tbx2b_*=16; Conover-Iman *posthoc* corrected p-values: wild type *vs. tbx2b* p=5.87 × 10^−5^). (**H**) Quantification of the fraction of M-opsin expressing S cones (double positive cells in B) in wild type, *tbx2b* mutants, and *tbx2a* mutants. *tbx2b* mutants have significantly GFP-positive S cones compared to wild type while *tbx2a* mutants do not. (Kruskal-Wallis H=21.101, p=2.61 × 10^−5^, n*_wild type_* = 58, n*_tbx2b_*=8, n*_tbx2b_*=8; Conover-Iman *posthoc* corrected p-values: wild type *vs. tbx2b* p=2.85 × 10^−6^, wild type *vs. tbx2a* p=1.0*, tbx2a vs. tbx2b* p=1.16 × 10^−3^). Bars represent averages, error bars correspond to standard deviations, and markers correspond to individual retinas.

### Tbx2a regulates M-opsin expression in L cones

To characterize the role of Tbx2a in the development of L cones, we used *tbx2b*^mcp1^ and *tbx2a*^mcp2^ mutants in combination with reporter lines for M opsin (*opn1mw2*:GFP) and for L cones (*thrb*:tdTomato) and quantified photoreceptor densities and homotypic NND (see STAR methods Key Resources Table). In wild-type larvae, M cones (GFP positive) and L cones (tdTomato positive) are regularly distributed with negligible overlap in expression between reporters (Figure 3A). We find that in both *tbx2* mutants, L cones (tdTomato-positive) are present in normal densities and their spacing is indistinguishable from wild type (Figure 3B-E). We also find the same phenotype described above, where cells that express M-opsin expression (GFP positive) have increased density and appear irregularly spaced (Figure 3B-C). In *tbx2b* mutants, L cones are not GFP positive and are likely normal, in agreement with the lack of expression of *tbx2b* in L cones. In contrast, the supernumerary GFP-positive cells in *tbx2a* mutants correspond to L cones (tdTomato and GFP positive), a sign of ectopic expression of M opsin in L cones (Figure 3D-F). By excluding the tdTomato and GFP positive L cones, we quantified the density and spacing of GFP-only cells, which we presumed correspond to M cones in *tbx2a* mutants, and again found that their density is not significantly different from wild type animals but there is a decrease in their nearest-neighbor distance, suggesting once more that M-cone arrangement but not number is impacted by the absence of UV cones (Figure 3G-H). These results suggest that, in *tbx2a* mutants, M cones are likely normal, but L cones lose the ability to repress M-cone fate, which erodes their subtype-specific identity.

**Figure 3:**
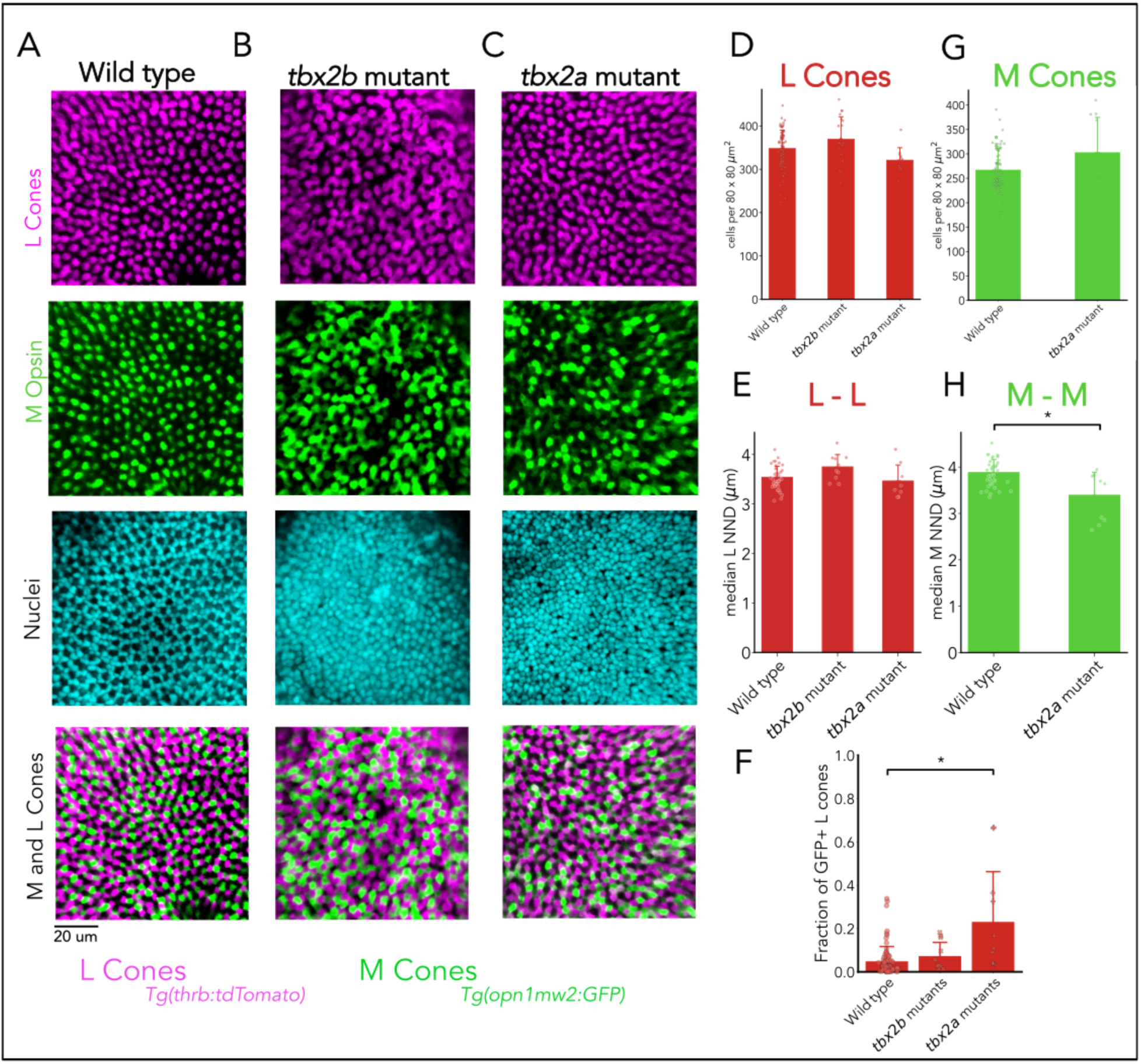
*tbx2a* represses M opsin in L cones. (**A-C**) Representative confocal images of the central retina of wild type (left), *tbx2b* mutants (center), and *tbx2a* mutants (right) at 5 dpf, in double transgenic larvae with labeled M cones (green), L cones (magenta), and nuclei (cyan). Both *tbx2b* and *tbx2a* mutants display an increase in GFP-positive cells. In *tbx2b* mutants the increase in GFP signal is excluded from tdTomato-positive cells, whereas *tbx2a* mutants have an increase in GFP signal that is restricted to tdTomato-positive cells. (**D-E**) Quantification of L and M cone numbers in wild-type, *tbx2a* and *tbx2b* mutant larvae. Compared to wild type, neither *tbx2b* or *tbx2a* mutants have significant changes in L and M cone densities (Kruskal-Wallis H=4.269, p=0.118, n*_wild type_* = 79, n*_tbx2b_*=10, n*_tbx2a_*=10). (**F-G**) Quantification of L and M cone spacing using median nearest-neighbor distances between homotypic photoreceptor subtypes. Both *tbx2b* and *tbx2a* mutants do not have significant changes in the spacing of L cones compared to wild type, *tbx2a* mutants do not have a significant decrease in spacing of M cones (Kruskal-Wallis H_L_=6.285, p=0.04376; Kruskal-Wallis H_M_ =21.156, p=2.54 × 10^−5^; n*_wild type_* = 40, n*_tbx2a_*=10, Conover-Iman posthoc corrected p-values: wild type vs. *tbx2a* p=0.00624) (**H**) Quantification of the fraction of GFP-positive L cones (double positive cells in C) reveals a significant increase only in *tbx2a* mutants (Kruskal-Wallis H=14.870, p=0.00059, n*_wild type_* = 80, n*_tbx2b_*=11, n*_tbx2a_*=11; Conover-Iman posthoc corrected p-values: wild type vs. *tbx2b* p = 0.190, wild type vs. *tbx2a* p=0.00055, *tbx2a vs. tbx2b* p=0.399). Bars represent averages, error bars correspond to standard deviations, and markers correspond to individual retinas.

### Tbx2 paralogs regulate expression of photoreceptor and subtype-specific genes

To validate and extend the phenotypes observed by imaging, we performed bulk RNA-sequencing on wild-type, *tbx2b-*mutant, and *tbx2a-*mutant heads at 5 days post fertilization (Figure S2A). Consistent with the loss of UV cones and increase in rods observed by imaging, *tbx2b* mutants exhibited significantly reduced expression of the UV opsin (*opn1sw1*) and increased expression of rhodopsin (*rho*). In contrast, *tbx2a* mutants showed significantly reduced UV-opsin expression but no change in rhodopsin levels, consistent with the incomplete loss of UV cones without changes in rods from imaging analyses. Moreover, comparisons between the two mutants reveal that UV-opsin expression is lower in *tbx2b* mutants than *tbx2a* mutants, again reflecting the complete *vs.* partial loss of UV cones. By assessing expression of S, M, and L opsins, we find a decreased expression of S opsin (*opn1sw2*) but only in *tbx2b* mutants (compared to wild type or to *tbx2a* mutants). M and L opsins have been duplicated and have differential expression across development.^30,31^ We find that, compared to wild type, the general developmental patterns are not altered in *tbx2* mutants, with low expression (at 5 days post fertilization) of *opn1mw3, opn1mw4,* and *opn1lw1* and higher expression of *opnmw1, opn1mw2,* and *opn1lw2* (Figure S2C). Yet, we see significantly decreased expression of *opn1mw2* in *tbx2a* mutants compared to wild type and *tbx2b* mutants, consistent with the paralog-dependent changes in M-opsin positive cells observed by imaging (compare Figure 2B-F and Figure 3C-F). In summary, these opsin expression changes mirror our imaging results, supporting our interpretation that *tbx2* loss alters photoreceptor subtype composition and subtype-restricted opsin programs.

To assess alterations in the developmental programs required for generation of photoreceptor subtypes, we assessed differences in the expression of relevant transcription factors. In zebrafish, rods can be generated through Nrl-dependent or Mafba-dependent activation of Nr2e3.^32^ While we detect no significant differences in *nrl* expression in either of the *tbx2* mutants, *nr2e3* expression is decreased in *tbx2a* mutants with a significant increase in *mafba (*Figure S2D), suggesting that these genes are still involved in the formation of normal rods. We find significant decrease of both *tbx2b* and *tbx2a* expression in both mutants, validating our mutations and consistent with the change of UV opsin and loss of UV cones. We find a significant decrease in *foxq2* expression in *tbx2b* mutants but not *tbx2a,* consistent with alterations in the development of S cones in *tbx2b* mutants. We also find a significant decrease in transcription factors important to generate M and L cones, particularly *thrb*, *six6a*, *six6b*, and *six7* (Figure S2D). The significant decrease in *thrb* in both mutants correlates with the significant decrease in *opn1lw2* expression (see above). To further quantify changes in rod versus cone-subtype differences, we assessed the expression of cone-enriched genes. Compared to wild type, we see reduced abundance of cone-enriched transcripts in both *tbx2* mutants including components of the phototransduction cascade and other markers of mature cone identity (Figure S2E-F). These transcriptional changes could be caused purely by the loss of UV cones or could also involve further changes in the other cone subtypes.

In summary we find that *tbx2b* mutants are unable to generate UV cones, switch the fate of these progenitors to become rods, and S cones have a hybrid S/M identity, while L cones and M cones appear to develop in normal numbers but with alterations in regulatory programs and opsin expression. In *tbx2a* mutants there is a partial loss of UV cones and L cones have a hybrid L/M identity while S cones, M cones and rods appear to develop in normal numbers, but also with alterations in regulatory programs and opsin expression. In other words, *tbx2* paralogs in zebrafish are key nodes in the regulatory programs that control the identity of all photoreceptor subtypes, with important roles for the generation of UV cones *vs.* rods and in the separation of identity between M *vs.* S and L cones. Given the multiple alterations in photoreceptor subtypes, we decided to test the functional consequences for vision, using assays for optokinetic response and prey capture.

### Luminance and motion detection are normal in *tbx2* mutants

To determine whether *tbx2* mutants can detect and respond to luminance and motion cues, we tested the optokinetic response (OKR) in immobilized 5 days post fertilization larvae, by quantifying eye movements evoked by black-and-white drifting gratings (spatial frequency: 0.016 cycles/°, speed: 60°/s) of low contrast (1 to 5%) (Figure 4A-C; see STAR Methods). We find that wild-type larvae have a normal OKR with slow tracking eye movements, interspersed with saccades, the fast eye movements that reset the eye position (Figure 4D).^33^ In wild-type larvae, eye velocity during tracking increases with contrast, with 1% contrast approaching the limit of detection (Figure 4E). We find no significant differences in OKR performance between wild type, *tbx2b* mutants, and *tbx2a* mutants across the contrast range tested (Figure 4E), supporting that luminance- and motion-driven visual responses are preserved in both mutants. Since larval zebrafish OKR depends primarily on pooled input from L and M cones,^34^ these results suggest that these cone subtypes remain functional and correctly wired with visual circuits in both mutants, despite the hybrid L/M-cone identity observed in *tbx2a* mutants, and the alterations in developmental programs and in L- and M-opsin expression.

**Figure 4:**
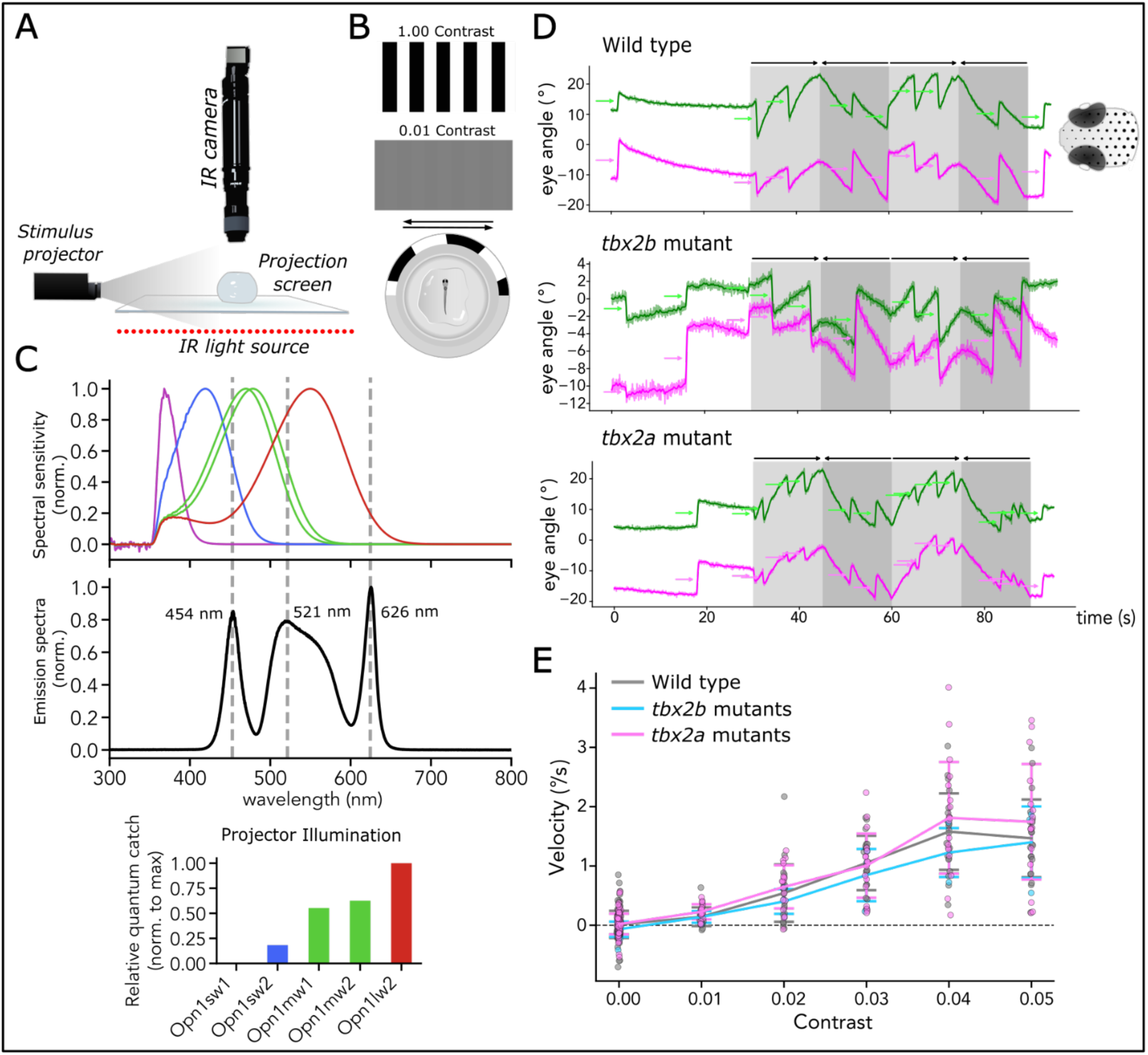
Luminance and motion detection are normal in *tbx2* mutants. A) Schematic of optokinetic response (OKR) recording setup. Individual immobilized 5 dpf larvae were recorded from above with an IR camera while a black-and-white grating was projected onto an acrylic screen surrounding the immobilized zebrafish (B) Example stills of the OKR stimulus at 100% and 1% contrast (top). Arrows indicate stimulus direction and reversal every 15 s (middle). Enlarged top view of a larva embedded in methylcellulose (bottom). (C) Spectral sensitivity curves for cone opsins expressed at this developmental stage (corrected by lens absorption), including Opn1sw1, Opn1sw2, Opn1mw2, Opn1mw1 and Opn1lw2 (top), measured emission spectra from the projector (middle), and calculated relative quantum catches for the indicated cone opsins (bottom). (D) Example left- and right-eye angle traces (left in pink and right in green) at 5% contrast from wild-type, *tbx2b* mutant, and *tbx2a* mutant larvae. Saccades appear as sharp deflections and are indicated by arrows, whereas slow tracking occurs between saccades. (E) Average eve velocities during slow tracking of the stimulus in wild type, *tbx2b* mutant, and *tbx2a* mutant larvae across tested contrasts (1 - 5%). Higher contrast elicits faster tracking, with 1% contrast approaching the limit of detectability. Average eye velocities increased significantly with contrast (x^2^(1) = 107.96, p = 2.74 x 10^-25) but did not differ by genotype (x^2^(1) = 0.41, p = 0.52; n_wt_ = 22, n*_tbx2b_* = 6, n*_tbx2a_* = 18), and there was no genotype-by-contrast interaction (x^2^(2) = 1.50, p = 0.22).

### Prey capture is disrupted in *tbx2b* mutants but preserved in *tbx2a mutants*

Larval zebrafish forage on unicellular organisms such as paramecia and rotifers. Successful prey detection and capture depend on UV-cone signaling, particularly from those in the ventro-temporal retina.^35,36^ To test whether complete or partial loss of UV cones impairs prey capture, we placed individual 7 days post fertilization larvae in a small arena with 40 rotifers in darkness or under red (632 nm), or UV (365 nm) illumination and quantified successful capture events over 4.5 min (Figure 5A-B; see STAR Methods for details). Wild-type larvae are unable to capture prey under dark or red illumination but readily detect and capture prey under UV illumination, confirming that our time-restricted assay requires vision and UV stimulation. While both *tbx2b* and *tbx2a* mutants failed to capture prey in darkness or under red illumination, *tbx2b* mutants, which have a complete loss of UV cones, also fail to capture prey under UV illumination, consistent with foraging deficits in *lor* mutants.^37^ Surprisingly, *tbx2a* mutants, which have a marked reduction in UV-cone densities, are still able to capture prey under UV illumination at rates comparable to wild type (Figure 5C). This suggests that the remaining UV cones in *tbx2a* mutants are functional and capable of supporting prey capture behavior. To further examine the hunting strategies, we performed frame-by-frame pose estimation in wild-type and *tbx2a* mutant larvae and found no significant difference in peak eye convergence prior to the strike, which is key for depth estimation and successful captures (Figure 5D; see STAR Methods).^38–42^ Together, these results show that UV-cone signaling underlies prey capture in larval zebrafish, and that even a reduced UV cone population, as seen in *tbx2a* mutants, is sufficient to support this behavior, in spite of the signaling constraints.^35^

**Figure 5:**
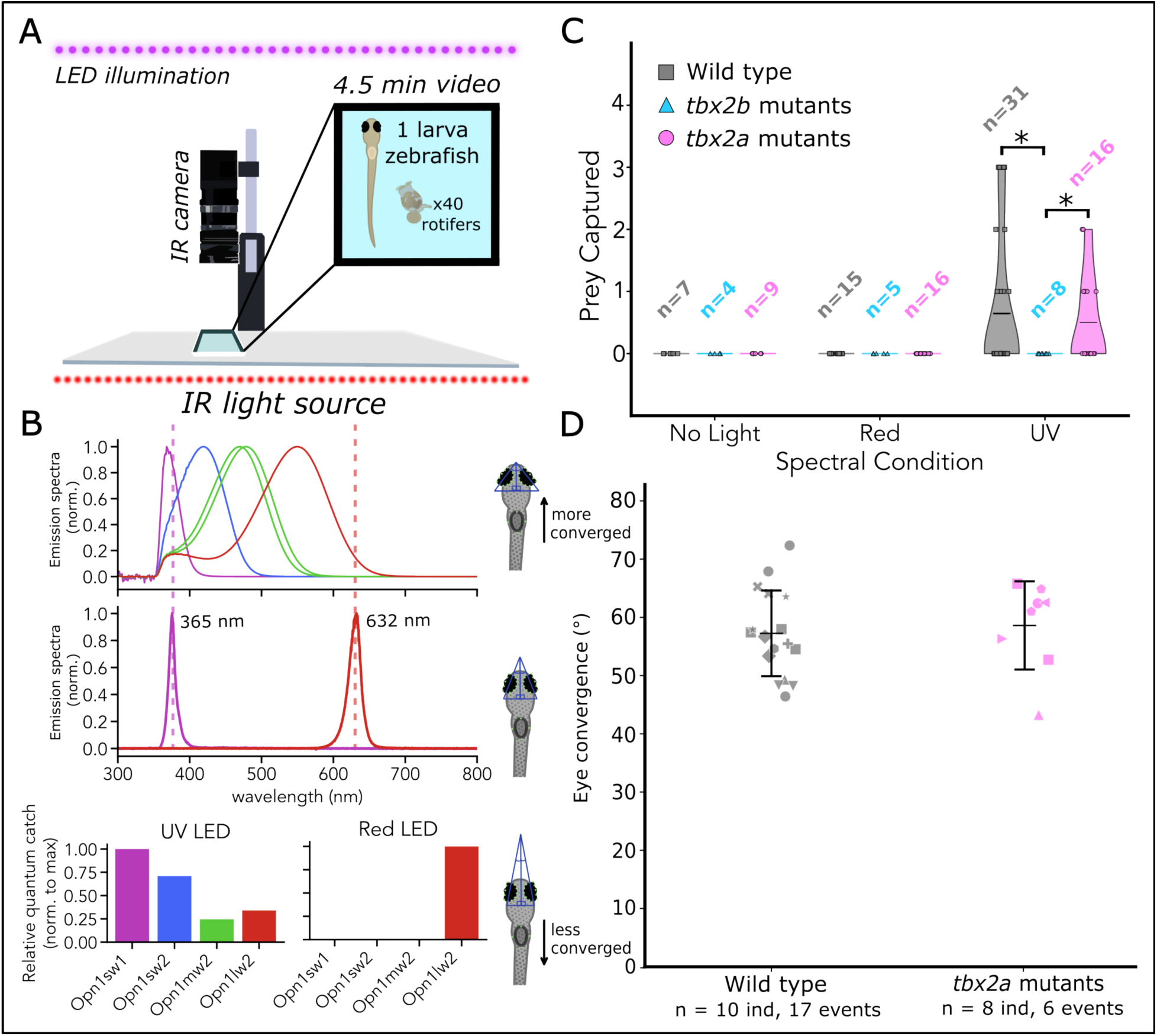
Luminance and motion detection are normal in *tbx2* mutants while prey capture is disrupted in *tbx2b* mutants but preserved in *tbx2a* mutants. (A) Schematic of the prey-capture setup. Individual 7 dpf larvae were recorded with an infrared (IR) camera with IR illumination from below and visible LED illumination from above. The enlarged chamber view depicts one larva and a representative rotifer (not to scale); 40 rotifers were added in each 4.5 min trial. (B) Spectral sensitivity curves for cone opsins expressed at this developmental stage (corrected by lens absorption), including Opn1sw1, Opn1sw2, Opn1mw1, Opn1mw2 and Opn1lw2 (top), measured emission spectra of the LEDs used in prey-capture trials (middle), and calculated relative quantum catches for the indicated cone opsins under each illumination condition (bottom). (C) Number of prey captured in wild-type, *tbx2b* mutant, and *tbx2a* mutant larvae at 7 dpf under no-light, red, and UV conditions. Each point represents one larva; n denotes the number of larvae tested in that illumination condition. Violin plots estimate the distribution of captures, and black lines indicate the mean. Under UV illumination, there was no significant difference across genotypes was not significant (Kruskal-Wallis: H = 3.470, p = 0.1764). Pairwise comparisons show significantly reduced capture rates in *tbx2b* mutants versus wild type (Mann-Whitney U, one-sided, p = 0.0449) and versus *tbx2a* mutants (Mann-Whitney U, one-sided, p = 0.0287) under UV light. Under no-light and red-light conditions, all groups captured zero prey. (D) Maximal eye convergence angles in wild type and *tbx2a* mutant larvae in successful captures (n_wt_ = 10, n*_tbx2a_* = 8); *tbx2b* mutants were not included as they did not capture any prey. Each point represents one successful capture; individual larvae are marked by the same symbol. Horizontal bars indicate group mean ± SD. We did not find significant differences (likelihood-ratio test p = 0.658; Wald p = 0.675; estimated genotype effect = +1.37°, 95% Cl [-5.05°, 7.80°]).

## DISCUSSION

A central challenge in developmental biology is understanding how transcription factors play key roles to first make closely related fates distinct during specification and to maintain these identities as separate throughout the lifespan of an organism. Here we show that the duplicated T-box genes *tbx2a* and *tbx2b* have subfunctionalized to meet both demands in the zebrafish retina. Loss of *tbx2* disrupts UV-cone generation with consequences for rod development, but also destabilizes the maintenance of cone subtype boundaries, leading to ectopic activation of M-cone programs in S and L cones. Together, these results argue that *tbx2* paralogs act as key regulatory nodes that coordinate fate allocation and identity maintenance, with selective consequences for visually-guided behavior.

### Tbx2 is a central regulator of all photoreceptor subtypes

Decades of research have underscored the importance of transcription factors to control cell-fate decisions in photoreceptors.^43,44^ Our results demonstrate that *tbx2b* and *tbx2a* are critical for not only UV cone development, but to maintain the cell identity of other photoreceptor subtypes. Our *tbx2* mutants also highlight that a single transcription factor can have multiple functions through their expression in different cell subtypes. UV-cone precursors utilize *tbx2b* to repress rod-specific genes and express both *tbx2b* and *tbx2a* to activate UV cone-specific genes. At the same time, S and L cones express *tbx2b* and *tbx2a* respectively to repress M-cone fate—particularly expression of the *opn1mw2* opsin. Given the differences in phenotypes, it is likely that the *tbx2* paralogs act both in shared and independent regulatory networks, which will be important to determine. For example, T-Box transcription factors co-regulate each other to create patterning differences across the chicken retina.^45^ This is also supported in our transcriptomics data: *tbx1* is upregulated in both *tbx2* mutants, while *tbx2*, *tbx3a, tbx5a, tbx6,* and *tbx16* are significantly reduced in both mutants compared to wild type. In addition, each *tbx2* paralog is downregulated in both mutants, suggesting a model where the two *tbx2* genes may be co-regulated, lending to mechanisms to control identity over all five photoreceptor subtypes.

### Tbx2 is enriched in UV cones across vertebrates

Much of the work on transcriptional regulation of photoreceptors has been in rods, while the transcription factors that govern cone fates are less understood. In mice, OTX2 promotes photoreceptor precursor identity,^43,46^ NRL drives rod fate,^47–49^ while THRB promotes L-cone fate.^50,51^ Precursors that do not express either NRL or THRB generate short-wavelength sensitive cones,^50^ leading to the idea that this is the default identity that is repressed by NRL and THRB. Across vertebrates, however, short-wavelength cone lineages (UV cones in zebrafish and many non-mammalian species; S cones in mammals and primates) show a conserved transcriptional signature: single-cell and bulk transcriptomic datasets consistently report specific TBX2 expression in *opn1sw1-*expressing cones in human,^52^ primate,^6,53^ mice,^54^ zebrafish^12^ and chicken,^6,55,56^ suggesting that TBX2 is important for the development of the short-wavelength cone lineage across vertebrates. Our functional data in zebrafish support a model where generation of the short-wavelength sensitive cones is not a passive “*default*” route, but rather requires active transcriptional regulation, controlled by TBX2, to activate subtype-specific programs and restrict alternative photoreceptor fates.

### Heterogeneity amongst a single photoreceptor subtype

In contrast to the rod-dominated mouse model, evidence in zebrafish suggest that cones and rods both require activation of specific transcription factors to assume final cellular identity. In larvae, *tbx2b* expression is required to repress rod-specific genes in order to generate UV cones, if absent all UV cones become rods (Figure 1G-K).^24,25^ Since UV cones are still present in *tbx2a* mutants, our results suggest at least two pathways for UV-cone generation in zebrafish: one pathway depends on both Tbx2b and Tbx2a, while in the other Tbx2a is dispensable. Determining the exact downstream mechanisms for how each of these transcription factors control photoreceptor diversity will be critical to fully understand how UV-cone identity is determined and how novel cell subtypes emerge during evolution. Interestingly, zebrafish L cones also have distinct subpopulations that can be revealed, amongst other markers, by tracking differences in the expression of the two L opsins (*opn1lw1* and *opn1lw2*). Each of the L opsins is differentially expressed across development (*opn1lw2* expression is higher in larval stages) and then becomes regionalized to the dorsal and central retina.^30,31^ Like Tbx2, Thrb plays a similar dual role, being involved in the generation of L cones and in controlling the differences between the two subpopulations.^57–59^

### Tbx2 paralogs stabilize cone subtype identity and prevent hybrid fates

Our *tbx2* mutants illustrate that developing cells acquire their final subtype-specific identity in stages, where transcription factors play distinct roles. In *tbx2b* mutants, the inability of progenitors to become UV cones is accompanied by a UV-to-rod fate shift, suggesting that Tbx2b is involved in an “early” fate decision. The lack of fate shift in *tbx2a* mutants indicates that Tbx2a acts differently and likely involved in a subsequent fate decision and reinforces that Tbx2b is the main repressor of this fate switch. Switches in fate have also been documented in *thrb* and *nrl* mutants, where inability of progenitors to become L cones or rods, respectively, forces a switch to a UV cone fate.^60,61^

In comparison, S and L cones are, initially, generated appropriately in the *tbx2* mutants (normal densities and spacing), but subsequently fail to maintain their identity by ectopically expressing genes related to M cones. This suggests that S and L cones in *tbx2* mutants acquired a hybrid state between competing fates and that the Tbx2 paralogs act again as repressors, but at a later stage of photoreceptor development. These hybrid photoreceptor states have also been observed in *samd7* mutants, where L cones abnormally co-express L and UV opsins,^60^ indicating that repressing related fates throughout the lifespan of a cell is a general mechanism for active maintenance of cell identity and function.^2,62^ Interestingly, other species seem to have co-opted such hybrid states for ecological advantages. In mice, L cones co-express L and UV opsins, following a dorso-ventral gradient that matches the spectral constraints of their visual environment.^63,64^ In cichlids, opsin expression in photoreceptors is plastic and hybrid states are common as opsin expression switches during development or in response to environmental cues.^65–67^ In the future, it will be interesting to understand the regulatory networks that allow for the maintenance and/or dynamic control of these hybrid states. Our transcriptomic data indicates that Tbx2 may be key to regulate such opsin-expression programs, but its influence expands to the control of other transcriptional regulators and of genes involved in phototransduction, supporting that *tbx2* activity contributes to stabilizing subtype-specific gene-regulatory networks beyond opsin choice.

### OKR remains intact in *tbx2* mutants despite altered cone identity

The optokinetic response (OKR) is widely used, clinically in humans and experimentally in many vertebrates, to detect visual deficits ranging from blindness to disruption of specific signaling pathways. In zebrafish, the OKR is driven by the combined input from L and M cones, with little contribution from short-wavelength sensitive cones (S and UV).^34,68,69^ Thus, mutants lacking functional L cones lose OKR responses to long-wavelength stimuli.^59^ Given the alteration in L-cone fate in *tbx2a* mutants (Figure 3) and the decrease in L-opsin expression in both mutants (Figure S2C), we decided to test OKR. We found that *tbx2* mutants retain robust luminance-driven motion responses across decreasing contrasts, which suggests that L and/or M cones are still providing the visual input required for this reflex and that the downstream motion circuitry remains intact.

### UV-cone loss reveals distinct consequences for prey capture in *tbx2* mutants

We used prey capture to assess the behavioral consequences of developmental alterations in UV cones^35,70–72^. We find that *tbx2b* mutants, which completely lack UV cones, are unable to capture any prey using vision, consistent with prior work on *lor* mutants and UV-cone ablation^35,71^. This finding links *tbx2b*-dependent UV-cone fate to prey-capture behavior and shows that developmental programs establishing photoreceptor subtype identity can selectively shape visual function, and that the specificity of this task cannot be compensated by circuit rearrangements. By contrast, the preserved prey-capture performance of *tbx2a* mutants provides a more nuanced view of this relationship. Despite losing about a third of their UV cones, which we expected to impair prey detection, *tbx2a* mutants captured prey at rates comparable to wild type under UV illumination. Optical constraints, UV-cone densities, and prey size predict that a single UV cone is likely to mediate the initial prey detection.^35^ A reduced UV-cone population should still impair capture by degrading the ability to make binocular depth estimation as larvae approach the prey.^73^ Further studies will be required to understand how these visual computations are achieved normally or in the face of degraded visual input faced by *tbx2* mutants. Particularly, it will be important to understand changes in the retinal circuit that may provide functional compensation. For example, it has been described that H3 horizontal cells, which normally connect abundantly with UV cones, increase their connectivity with S cones in *lor* mutants.^74^ It is likely that other retinal circuits are rearranged as the identity of photoreceptor subtypes is altered; the nature of these changes is likely to dictate the differences in prey capture we describe between *tbx2* mutants.

## ACKNOWLEDGMENTS

Research reported in this publication was supported by the National Eye Institute Pathway to Independence Award (K99EY030144-01, R00EY030144 JA) and by the National Eye Institute Ruth L. Kirschstein Predoctoral Individual National Research Service Award (F31EY038082 CMH). The content of this manuscript is solely the responsibility of the authors and does not necessarily represent the official views of the National Institutes of Health.

## AUTHOR CONTRIBUTIONS

Conceptualization, J.A., C.M.H., and A.S.L.; methodology, J.A., C.M.H., A.S.L. and S.A.; Investigation, J.A., C.M.H., A.S.L., S.A., and A.K.R.; Writing – original draft, J.A., C.M.H., A.S.L., and S.A.; Writing – review & editing, J.A., C.M.H, and A.S.L.; Funding acquisition, J.A. and C.M.H.; Resources, J.A.; Supervision, J.A.

## DECLARATION OF INTERESTS

No competing interests declared

## DECLARATION OF GENERATIVE AI AND AI-ASSISTED TECHNOLOGIES IN THE WRITING PROCESS

During the preparation of this work, the authors used University of Maryland-owned chatbot - TerpAI (terpai.umd.edu) to edit minor organization in introductory paragraphs. After using this tool or service, the authors reviewed and edited the content as needed and take full responsibility for the content of the publication.

## KEY RESOURCES TABLE

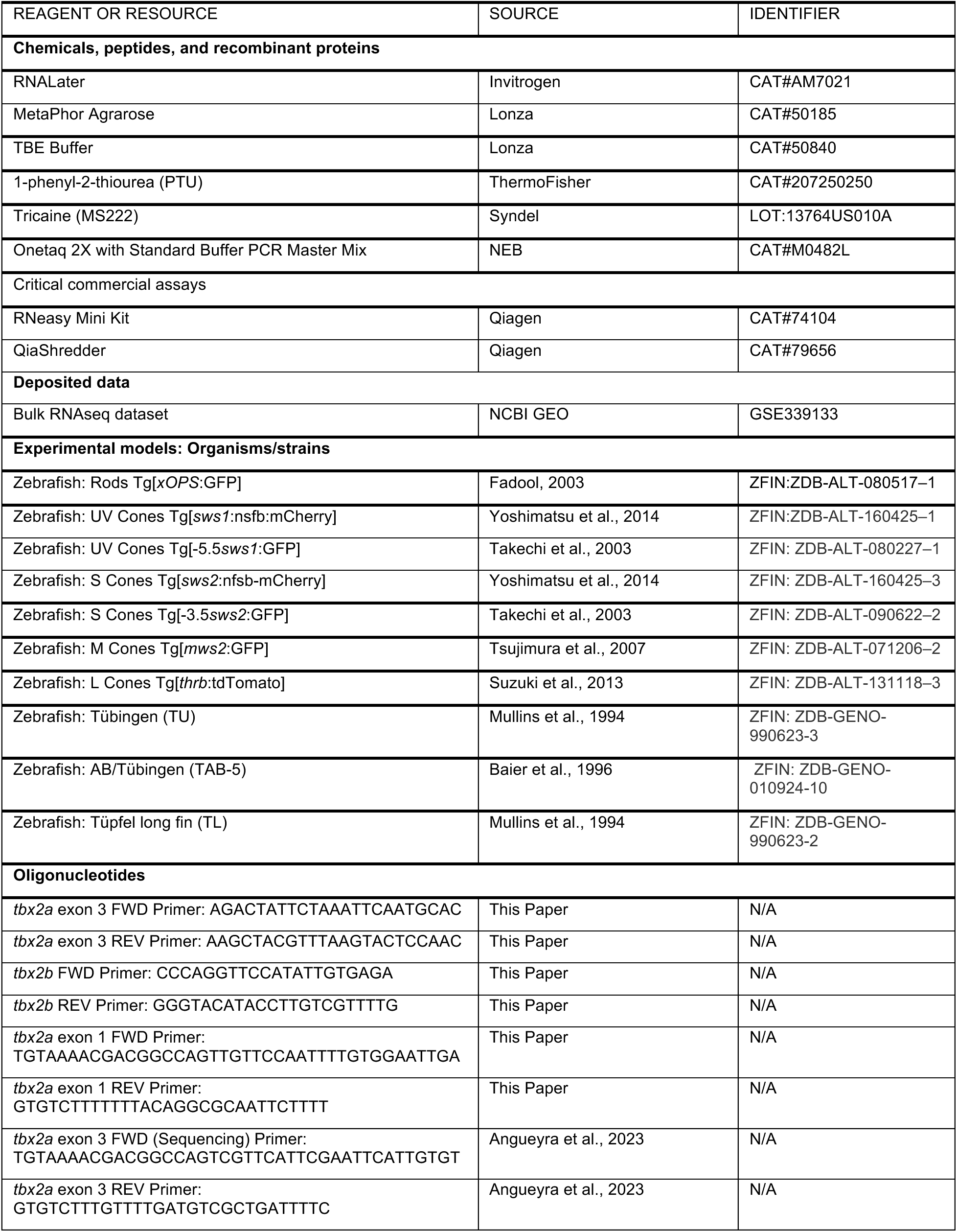
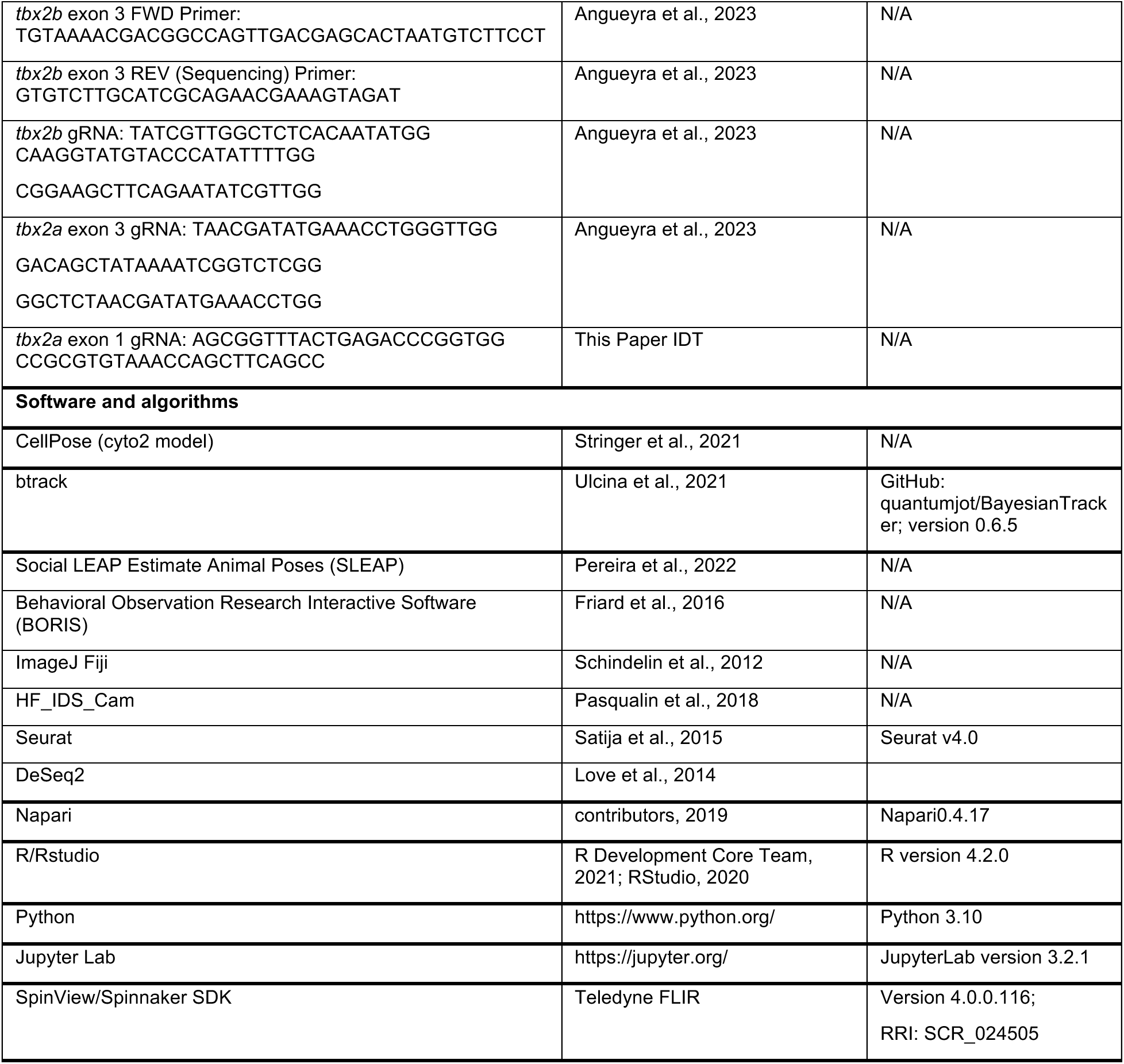

## EXPERIMENTAL MODEL AND STUDY PARTICIPANT DETAILS

### Animals

We grew zebrafish larvae at 28°C in E3 embryo media (5 mM NaCl, 0.17 mM KCl, 0.33 mM CaCl2, and 0.33 mM MgSO4, pH = 7.2) under a 14 h:10 h light-dark cycle (lights on from 8 A.M. to 10 P.M.). Before 20 hours per fertilization, we added 0.003% 1-phenyl-2-thiourea (PTU) to the embryo medium to block melanogenesis. All work performed at the University of Maryland and was approved by the UMD Animal Use Committee under animal study protocol #R-MAR-26-10. For all work with mutants, larvae were examined at 5 days post fertilization. At this age, sex cannot be predicted or determined, and therefore the sex of the animals was not considered. The transgenic lines used in this study are listed in Key Resources Table.

### Generation of *tbx2* Mutants

To create loss-of-function mutations in the *tbx2* paralogs, we used CRISPR-based mutagenesis by injecting RNP complexes. For each gene, we combined 3 different guides, all targeting the DNA-binding domain located in exon 3 (key resources table). We grew clutches with mutant embryos to sexual maturity and outcrossed to isolate F1 heterozygous founders. We outcrossed a single heterozygous F1 founder to generate a colony of F2 heterozygous individuals and then intercrossed to yield homozygous mutants. To identify the mutation, we isolated genomic DNA from the bodies of 5 days post fertilization larvae and PCR-amplified the genomic region containing the mutation (see key resources table) and used Sanger sequencing, to identify an 8 bp deletion in *tbx2b^mcp1^*mutants (chr15:29,853,882-29,853,889 [GRCz12/danRer12]) and a 14 bp deletion and 27bp insertion in *tbx2a^mcp2^* mutants (chr5:61,843,488-61,843,417 ([GRCz12tu/danRer12]), both predicted to result in a frameshift and a truncated protein product that lacks critical residues in the DNA-binding domain (Figure 1). These mutants are referred to throughout the paper as *tbx2a* mutants and *tbx2b* mutants. Subsequent genotyping was performed as described in “Genotyping”. To create a second mutation in the *tbx2a* gene for complementation studies, we combined 2 different guides, both targeting the first exon of *tbx2a* (key resources table), and followed the same protocol. We identified a 7 bp deletion also predicted to result in a frameshift and a truncated protein product that lacks the entire DNA-binding domain.

## METHOD DETAILS

### Sample Preparation and Imaging

For larval imaging, we enucleated eyes from fixed larvae using electrically-sharpened tungsten wires.^75^ We placed isolated eyes on a coverslip and oriented photoreceptors facing the coverslip before using a small drop of 1.0% low-melting point agarose to fix them in place. Upon solidification, we added a setting mounting medium (10% polyvinyl alcohol type II, 2.5% DABCO, 5% glycerol 25 mM, Tris buffer pH 8.7 and 0.5 μg/mL DAPI) and placed the coverslip on a glass slide, separated by a spacer (double-sided duct tape) to avoid compression. We used the bodies of the larvae for genotyping and imaged the corresponding larval retinas using a Nikon AxR resonant-scanning confocal microscope with a 25 x, 1.10 NA water-immersion objective and and high-sensitivity GaAsP detector units. We acquired z-stack images from a 80 µm x 80 µm square area of the central retina (dorsal to optic nerve) for photoreceptor quantification every 0.4–0.5 µm with a pinhole of 1.2 Airy Unit (AU) at a 2048 × 2048 pixel resolution. We excited fluorophores using diode/solid-state laser lines at 405 nm (DAPI), 488 nm (eGFP), and 561 nm (mCherry and tdTomato) and detected the emission on GaAsP channels using the following collection windows: DAPI 429–474 nm, eGFP 499–528 nm, mCherry 592–625 nm, and tdTomato 592–721 nm. To route the excitation light to the scan head, the instrument’s internal beam-combining optics are directed to the specimen using a multi-band main dichroic mirror. To eliminate bleed-through emission from other fluorophores, we separated DAPI and eGFP emission spectra into detection channels using a secondary dichroic mirror/filter set prior to detection by GaAsP photomultiplier tubes. We obtained lateral views of 5 dpf larvae using an Axiocam 208 color camera on a Zeiss Axio Zoom.V16 microscope or a Leica FLEXACAM C1 color camera on a Leica M165FC stereomicroscope (Figure S1I).

### Genotyping

We extracted DNA from the bodies of larvae at 5 days post fertilization using the HOTSHOT method.^76^ We placed samples in 25 µL of 25 mM NaOH with 0.2 mM EDTA, heated to 95 °C for 30 min, and cooled to 4 °C. Then we neutralized the solution by adding 25 µL of 40 mM Tris-HCl and vortexed the samples. For genotyping of samples in Figure 1, we used a fluorescent PCR method.^77^ We added the M13F adapter sequence to forward primers and the PIG-tail adapter sequence to reverse primers (see Key Resource Table) and used incorporation of fluorescent M13F-6FAM for detection.^77^ For all other samples (Fig. 2-4), we used non-fluorescent PCR and a high-resolution gel electrophoresis method.^78^ We PCR amplified and used 3.5 – 4% high-resolution MetaPhor Agarose gel in TBE buffer (Lonza) to visualize respective −8 bp and +13 bp size differences between *tbx2* mutants and their respective heterozygous and wild-type siblings. The PCR mixture (1 x), for a 13 µL reaction, contained forward primer (0.158 µM, 0.25 µL), reverse primer (0.316 µM, 0.25 µL), Onetaq 2X with Standard Buffer PCR Master Mix (NEB) (6.25 µL), water (3.75 µL), and 2.5 µL of DNA. We used the following PCR protocol: (1) 94 °C denaturation for 30 s, (2) 34 cycles of 94 °C for 15 s, 45 – 68°C for 30 s, 68 °C for 30 s (3) final extension at 72 °C for 10 min, (4) hold at 22°C. All primers and expected sizes are provided in Key Resource Table.

### Sample preparation for Bulk RNA-Sequencing

Since *tbx2a* mutants could grow to adulthood, we intercrossed *tbx2a^mcp2^*/ *tbx2a^mcp2^* mutants to produce all *tbx2a* mutant broods. For wild type samples, fish with no mutations were intercrossed from transgenic *Tg[mws2:GFP]* tanks. Since *tbx2b* mutants had low survival rates, we could not grow robust numbers to interbreed full mutants. To generate *tbx2b^mcp1^*/ *tbx2b^mcp1^* mutants, we intercrossed *tbx2b^mcp1^* heterozygotes to produce mixed genotype offspring. At 5 days post fertilization, we euthanized larvae by tricaine overdose, dissected the head of the larvae and stored them in RNAlater (Invitrogen) at −80°C; corresponding bodies were collected for genotyping. For each replicate we pooled 15 - 22 heads and obtained four wild-type, three *tbx2b^mcp1^*-mutant, four and *tbx2a^mcp2^*-mutant replicates. We homogenized head tissue using the Qiashredder columns (Qiagen) and extracted RNA using the RNeasy Micro Kit (Qiagen). Purified RNA was shipped on dry ice for 3’ end RNA-seq using a commercial service (Plasmidosaurus), converting mRNA to complementary DNA (cDNA) libraries through reverse transcription using a poly(dT)VN primer and second-strand synthesis, followed by tagmentation, library indexing, and amplification. To capture differential gene expression, cDNA libraries were used to perform 3’ end sequencing, with a read length of 90 bp.

### Bulk RNA Sequencing Analysis

We converted raw sequencing into FastQ files and performed demultiplexing with BCL Convert v4.3.6 and fqtk v0.3.1. All subsequent FastQ files were filtered and trimmed using FastP v0.24.0 with a minimum Phred quality score of 15, and minimum length requirement of 50 bp. We then aligned reads to the appropriate reference genome using STAR aligner v2.7.* with non-canonical splice junction removal and output of unmapped reads. In order to sort the BAM files, we indexed using samtools v1.21. To eliminate artificial, duplicate copies of genetic sequences generated during Next-Generation Sequencing (NGS), we ran UMICollapse v1.1.0. For quality control, we mapped reads with alignment quality metrics, strand specificity, and read distribution across genomic features using RustQC v0.2.1 and generated a comprehensive Quality Control (QC) report using MultiQC v1.33. To measure RNA levels, we used featureCounts (subread package v2.1.1) with strand-specific counting, multi-mapping read fractional assignment, exons and three prime UTR as the feature identifiers, and grouped by genotypes (wild type, *tbx2b* mutants, *tbx2a* mutants). We annotated final gene counts with gene biotype and other metadata extracted from the reference GTF file. Once final counts were established, we calculated sample-sample correlations for sample-sample heatmap and PCA on normalized counts (TMM, trimmed mean of M-values) using Pearson correlation. To identify genes or transcripts that are significantly upregulated or downregulated between genotypes, we performed differential expression analysis using Deseq2 and pcaExplorer for initial visualizations.^79,80^ Genes were considered as differentially expressed if fold-enrichment >1.5, p-value <0.01 and the estimated false positive rate or p-adjusted <0.1. In addition, genes were required to have positive reads in >50% of the enriched samples.

### Image Analysis

We imported confocal z-stacks of the central region of the retina (80 µm x 80 µm) into Napari.^81^ We created maximum-intensity projections using a small subset of the z-stack (2–10 planes) that ensured that we captured all photoreceptor cells in the region into a single image. We then used Cellpose, to segment photoreceptors in each image, using the cyto2 model.^82^ Finally, we manually corrected the segmentation to ensure all photoreceptors were properly counted. To quantify cone spacing, we used the conventional nearest neighbor distance (NND).^14^ We estimated the centroid of photoreceptors and calculated euclidean NND for each photoreceptor and plotted the median for each individual. To report reliable spacing and density counts of M cones, we used the M-cone segmentation (GFP-positive, mCherry-negative) from the cross labelling of S and M cones for *tbx2b* mutants and M-cone segmentation (GFP-positive, tdTomato-negative) from the cross labelling of M and L cones in *tbx2a* mutants. To quantify the GFP signal in S and L cones, we imaged Tg(*thrb*:tdTomato) or Tg(*opn1sw2*:nfsB-mCherry) and segmented the red channel and used it to create masks for the green channel. We normalized the GFP signal across each image and used a 27-pixel erosion (to avoid effects due to optical blurring) before calculating the average normalized GFP signal contained within each S cone or L cone. In the original work that established the Tg(*opn1mws2*:GFP) line, it was noted that a subset of S cones in control larvae are GFP-positive.^29^ We were able to identify these cells using a GFP signal threshold of 0.265 (4.987% of control S cones), and again used this same threshold to quantify the fraction of GFP-positive S cones in both control and each *tbx2* mutant larvae. Subsequently, to quantify M cone densities in these mutants, we performed manual counts of single positive cells (‘GFP only’) by excluding cells previously segmented as L or S cones. By plotting the distribution of GFP signal in L cones, we were able to establish a threshold of 0.205 that was exceeded by only 4.86% of L cones in wild-type larvae and used it to classify L cones as GFP positive in both control and mutant larvae.

### OKR testing

We performed optokinetic response (OKR) assays on 5 days post fertilization larvae, including wild type (n = 22), *tbx2b* mutants (n = 6), and *tbx2a* mutants (n = 18). All larvae were tested under binocular viewing conditions between 12:00 pm and 6:00 pm. We immobilized individual larvae in an upright position in 3% methylcellulose on the lid of a 35 mm petri dish. We recorded videos using an infrared-sensitive monochrome camera (IDS UI-3360CP-NIR-GL Rev.2) fitted with a 900 nm long-pass infrared filter (FEL0900, Thorlabs) and illuminated from below with 940 nm infrared LEDs repurposed from commercially available home-security cameras. Our visual stimuli consisted of vertically oriented square-wave gratings with a spatial frequency of 0.016 cycles/°, and velocity of 60°/s. We projected the stimuli with a digital projector (Texas Instruments LCR 4500) onto a cylindrical screen made from an acrylic lightbulb that surrounded the larva. The screen was 30 mm from the zebrafish eye. We recorded videos at 25 frames per second (25 fps)—using the HF_IDS_Cam plugin for image J—that included 30 s without stimulation (0% contrast) and 60 s with stimulation.^83^ To prevent desensitization, we switched the direction of the stimulus every 15 s. Single larvae were tested at multiple contrasts in descending order (1%, 2%, 3%, 4%, or 5%), with at least one minute of rest between trials. After behavioral trials, we genotyped all larvae.

### Prey Capture Testing

We tested prey capture behavior on 7 days post fertilization larvae, including wild type (n = 53), *tbx2b* mutants (n = 17), and *tbx2a* mutants (n = 41). All larvae were tested between 9:00 am and 2:00 pm in one of three illumination conditions: no-light, red light (632 nm), or UV (365 nm). We performed prey-capture trials in custom-built 3.05 x 3.05 cm acrylic chambers (Figure 5A) with black sides and a clear bottom. We recorded individual larvae at 40 fps using a Blackfly S monochrome camera (BFS-U3-200S236M-C USB 3.1, Teledyne FLIR). The camera was fitted with a 780 nm long-pass infrared filter (FGL780M, Thorlabs) and illuminated from below with 800 nm infrared LEDs, while LED illumination from above corresponded to the assigned trial condition (UV, no-light, or red; Figure 5A) and recorded videos using SpinView 4.0.0.116 (Teledyne FLIR). Before each trial, we transferred individual larvae to the chamber and allowed them to acclimate for 5 minutes under the testing illumination. After acclimation, we began recording, introduced 40 rotifers to the chamber, and continued acquisition for 4.5 min. Immediately after the trial, we photographed the remaining rotifers and manually counted them to estimate prey consumption. After behavioral trials, we genotyped all larvae.

## QUANTIFICATION AND STATISTICAL ANALYSIS

### Imaging

We performed statistical analyses and data plots using Python in Jupyter notebooks.^84^ Values of data and error bars in figures correspond to averages and standard deviations, and for statistical comparisons we used Kruskal-Wallis tests with a p-value <0.01 required for significance, unless stated otherwise. For statistical comparisons in *tbx2* mutants, we performed Kruskal-Wallis tests on the three groups (wild type, *tbx2a* mutants and *tbx2b* mutants), and significant results were followed up with a post hoc Conover-Iman test with a Bonferroni adjustment of p-value.^85^ Samples sizes, test values and significance levels are stated in the figure legends. No randomization, blinding, or masking was used for our animal studies, and all replicates are biological.

### OKR

We used cellpose^82^ to segment the eyes of each larva across frames, and bTrack^86^ to bind left vs. right eyes through entire videos. We then fit ellipses to the segmented eyes to quantify eye angle and instantaneous velocity. For saccades, we identified events where velocity exceeded 95^th^ percentile and manually corrected any omitted or misidentified saccades. We also manually excluded non-OKR movements (*e.g.*, twitching) from analysis. Tracking of gratings was quantified by averaging the slow-phase velocity and taking into account the direction of stimulation. All identified saccades and tracking were binocular, allowing us to average traces between left and right eyes. To test if *tbx2* mutants have impaired OKR response at different contrast levels, we used a linear mixed model fitted by restricted maximum-likelihood estimation using the lmer procedure in the lme4 R package.^87^ We confirmed data normality through a Shapiro–Wilks test on residuals of the model. We extrapolated significance values using a Type II Wald χ2 test using the ANOVA function in the car package of R.^88^ Each individual per genotype was included in the model as random factors.

### Prey capture behavior

To quantify prey capture behavior, we reviewed each trial in BORIS (Behavioral Observation Research Interactive Software) and annotated successful prey-capture events.^89^ We used the number of successful capture events per larva during the 4.5 min trial for statistical analysis. Because our *a priori* hypothesis was that *tbx2b* mutants would show impaired prey capture under UV illumination we used one-sided Mann-Whitney U tests for comparisons involving tbx2b mutants. Under no-light and red-light conditions, all larvae captured zero prey, preventing any statistical testing (Figure 5C). An increase in eye convergence is a hallmark of active prey pursuits that typically occurs at the end of the approach phase preceding the strike, when larvae have to make binocular estimations of depth.^39,41,42^ We identified the time point and angle of greatest eye convergence working backwards from each identified capture using SLEAP^90^ to label approach pre-strike frames and estimate the long axis of each eye by fitting an ellipse using 12 labeled nodes per eye. The angle at the intersection of the two eye axes was taken as the convergence angle, with smaller values indicating greater convergence. Because some larvae had more than one successful prey-capture event, convergence angles were analyzed at the event level using a linear mixed-effects model with genotype as a fixed effect and larva identity as a random intercept (angle ∼ genotype + (1 | larva)), thereby accounting for repeated measurements from the same animal (Figure 5D).

### Measuring spectral sensitivity and emission in behavior assays

We measured the emission spectra of our behavioral stimuli with a FLAME spectrophotometer (FLAME-S-UV-VIS-ES, Ocean Optics) and QP1000-2-UV-VIS optical fiber (Ocean Optics).

For OKR, we measured emission spectra of the projector for white (red, green, and blue LED simultaneously on) (Figure 4D). For prey capture, we measured emission spectra from the red and UV LEDs (Figure 5B). We estimated relative cone spectral sensitivities using the Govadorvskii A1 visual pigment templates and the reported peak spectral sensitivities for each opsin.^91^ We corrected these curves using an estimated lens transmission function measured from adult fish. We calculated relative quantum catch for each cone type by integrating the dot product of the measured stimulus spectrum and the corresponding lens-corrected spectral sensitivity across wavelengths. We included larval cone opsins (*opn1sw1, opn1sw2, opn1mw2,* and *opn1lw2*), and excluded later-stage expressed opsins (*opn1mw3, opn1mw4, and opn1lw1*) (Figure 4D and 5B).^92^

---

**Supplemental Figure 1.**
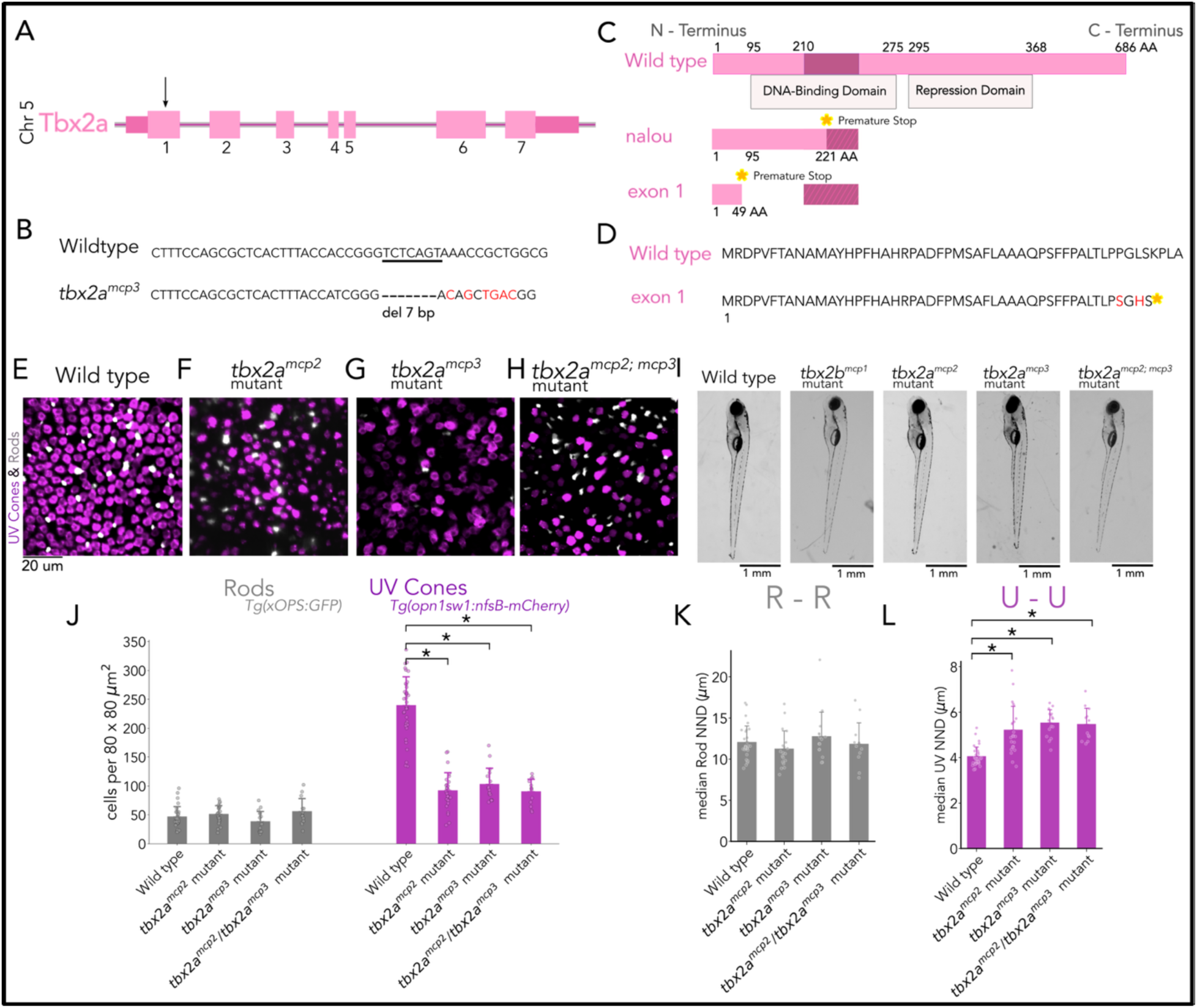
Complementation of two independent *tbx2a* mutations persist in partial loss of UV cones. (**A**) Map of *danio rerio tbx2a* gene structure (pink) on chromosome 5, arrow points to the first exon of the gene. (**B**) Mutation for exon 1 of *tbx2a* for wild type (top) and mutant (bottom) alleles. In *tbx2a^mcp3^* mutants, a 7 bp deletion (triangle) disrupts exon 1, encoding before the DNA-binding domain, therefore abolishing gene function. (**C**) Amino-acid diagram of *Danio rerio tbx2a* (pink) illustrates the primary sequence of the protein for wild type (top) and mutants (bottom). The amino acid positions that result from the third coding exon are shaded in darker pink for the protein. The highly conserved DNA-binding domain and less-well conserved repression domain is denoted by the shaded box underneath. Both mutations in exon 1 and 3 result in frameshifts that introduce a premature stop codon (indicated by yellow star). (**D)** Amino acid alignment of the first exon of wild type and *tbx2a^mcp3^* mutant the Tbx2a protein. (**E-H**) Representative confocal images of the central retina of wild type, *tbx2a^mcp2^*mutants, *tbx2a^mcp3^* mutants, and *tbx2a^mcp2/mcp3^* mutants at 5 dpf, in double transgenic larvae with labeled rods (grey) and UV cones (magenta). (**I**) Side profile of wild type, *tbx2b^mcp1^*mutants, *tbx2a^mcp2^* mutants, *tbx2a^mcp3^* mutants, and *tbx2a^mcp2/mcp3^* mutants at 5dpf. All mutations in *tbx2a* do not result in major malformations in larvae and appear physically similar to wild type. (**J**) Quantification of rods and UV-cone numbers in control, *tbx2a^mcp2^*and *tbx2a^mcp3^*, and *tbx2a^mcp2/mcp3^* mutant larvae. Bars represent averages, error bars correspond to standard deviations, and markers correspond to individual retinas. Compared to wild type, all *tbx2a* mutants have a significant decrease in UV cones (0.3860-fold for *tbx2a^mcp2^*and 0.4323-fold for *tbx2a^mcp3^*, 0.3786-fold for *tbx2a^mcp2/mcp3^*, Kruskal-Wallis H=62.993, p=1.34 × 10^−13^, n*_wild type_* = 36, n*_tbx2a_^mcp2^*=26, n*_tbx2a_^mcp3^*=16, n*_tbx2a_^mcp2 / mcp3^*=13; Conover-Iman posthoc corrected p-values: wild type vs. *tbx2a^mcp3^* p=1.03 × 10^−19^, wild type vs. *tbx2a^mcp3^* p=6.03, × 10^−14^, wild type vs. *tbx2a^mcp2/mcp3^*p = 6.08 × 10^−15^), while there was no significant difference between *tbx2a^mcp2^* and *tbx2a^mcp3^*, and *tbx2a^mcp2/mcp3^*double heterozygous mutants (*tbx2a^mcp2^* vs. *tbx2a^mcp3^* p=1.0, *tbx2a^mcp3^* vs. *tbx2a^mcp2/mcp3^*p=1). (**K-L**) Quantification of rod spacing in the central retina shows no significant changes in *tbx2a* mutants compared to wild type and to each other, whereas UV spacing shows significant changes of all *tbx2a* mutants compared to wild type. (Kruskal-Wallis H=0.547, p=1.0, n*_wild type_* = 56, n*_tbx2aexon3_*=8, n*_tbx2aexon1_*=9, n*_tbx2aexon3/1_*=9, Kruskal-Wallis H=44.796, p=1.02 × 10^−9^, n*_wild type_* = 33; Conover-Iman posthoc corrected p-values: wild type vs. *tbx2a^mcp2^* p=2.91 × 10^−8^, wild type vs. *tbx2a^mcp3^* p=4.55, × 10^−11^, wild type vs. *tbx2a^mcp2/mcp3^* intercross p = 4.55 × 10^−15^).

**Supplemental Figure 2.**
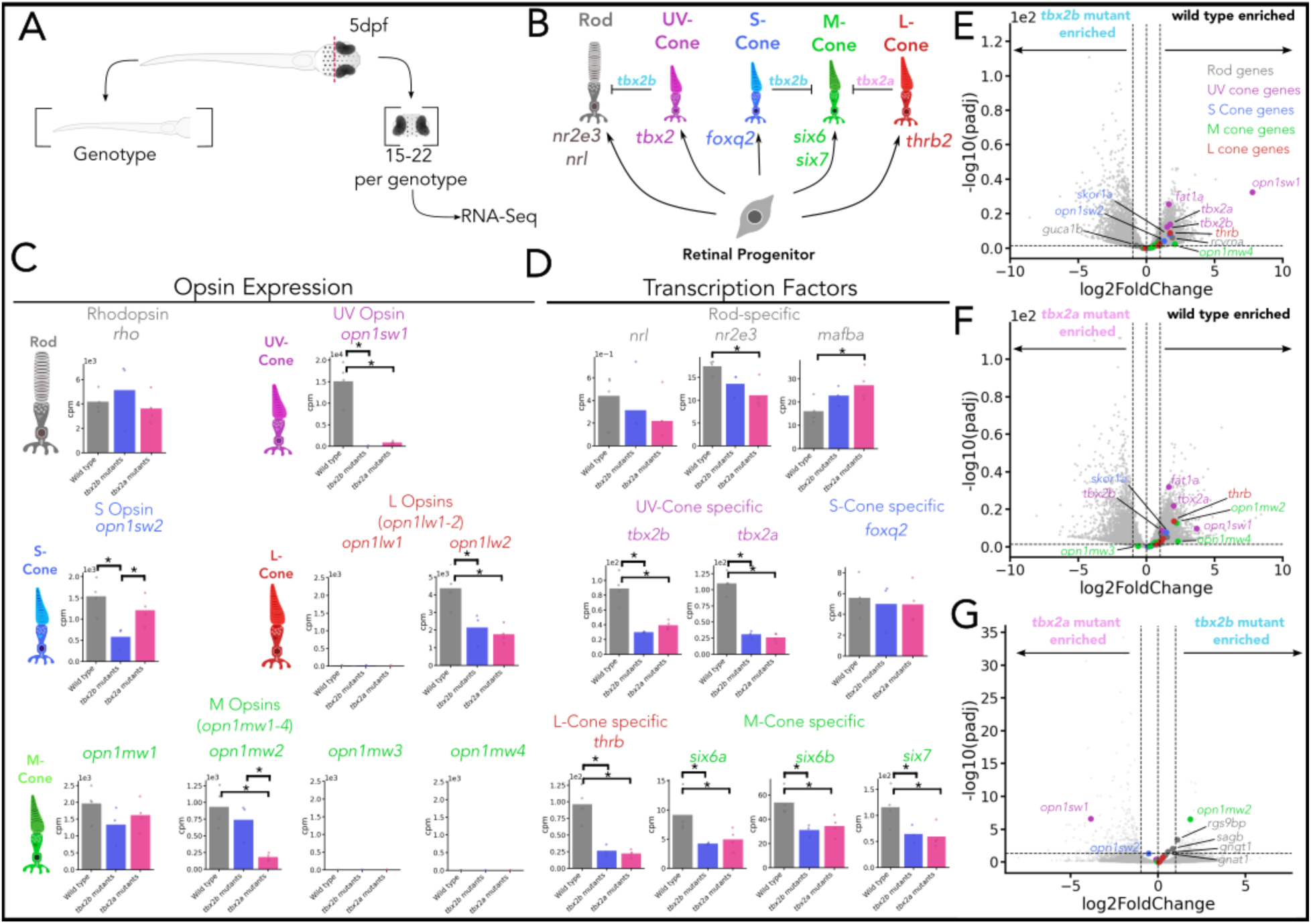
t*b*x2 mutants have alterations in the patterns of opsin expression, transcription factors that control photoreceptor identity and cone-related genes. (**A**) Diagram of sample prep. For each sample, 15-22 larval heads (5 dpf) were dissected, pooled and RNA was sequenced from wild type and each *tbx2* mutant (n*_wild type_* = 4, n*_tbx2b_^mcp1^*=3 n*_tbx2a_^mcp2^*=4). (**B**) Model for *tbx2* paralogs roles to create and maintain photoreceptor subtype-specific identities. Retinal progenitors (bottom) give rise to all photoreceptor subtypes: (gray) rod, (purple) UV cone, (blue) S cone, (green) M cone, and (red) L cone. Each arrow indicates a fate decision mediated by a particular transcription factor. Both *tbx2* genes are necessary for the generation of UV cones, whereas only *tbx2b* represses rod fate (left). In addition, both *tbx2* paralogs inhibit M-opsin expression; *tbx2b* inhibits M-opsin in S cones (middle), while *tbx2a* inhibits M-opsin in L cones (right). (**C**) Bar plots of opsin expression (counts per million after normalization for sequencing depth). The expression of UV opsin is significantly decreased in both *tbx2* mutants compared to wild type, while rhodopsin expression is significantly increased in just *tbx2b* mutants. S-opsin (blue) is significantly decreased in *tbx2b* mutants compared to wild type and *tbx2a* mutants (wild type vs. *tbx2b* mutant p=1.9 × 10^−5^, *tbx2b* mutant vs. *tbx2a* mutant p=0.0012). L-opsin (*opn1lw2*) is significantly decreased in both mutants compared to wild type (wild type vs. *tbx2a* mutant p=9.5 × 10^−15^, *tbx2b* mutant vs. *tbx2a* mutant p=1.14 × 10^−5^), and the larval-expressed M-opsin (*opn1mw2*) is significantly decreased in *tbx2a* mutants compared to both wild type and *tbx2b* mutants (wild type vs. *tbx2b* mutant p=1.9 × 10^−5^, *tbx2b* mutant vs. *tbx2a* mutant p=7.44 × 10^−7^). (**D**) Bar plots of photoreceptor-subtype specific transcription factor expression. Both *tbx2* paralogs are significantly downregulated in both mutants. Rod specific transcription factor, *nr2e3*, is significantly decreased only in *tbx2a* mutants compared to wild type, while *mafba* is only significantly increased in *tbx2a* mutants. All M-and L-cone specific transcription factors are significantly decreased in both *tbx2* mutants. (**E-G**) Volcano plots of bulk RNA-seq data comparing wild type to each *tbx2* mutant and to each other. Compared to both *tbx2b* and *tbx2a* mutants, wild type heads have enriched expression of cone specific genes. Whereas the *tbx2b* mutants have enriched rod specific gene expression compared to *tbx2a* mutants. We have highlighted a subset of genes known to have photoreceptor-subtype specific expression.

## Notes

### Competing Interest Statement

The authors have declared no competing interest.

